# Do identification guides hold the key to species misclassification by citizen scientists?

**DOI:** 10.1101/2025.04.10.648156

**Authors:** Fergus J Chadwick, Daniel T Haydon, Dirk Humseier, Jason Matthiopoulos, Otso Ovaskainen

## Abstract

1. Citizen science data often contain high levels of species misclassification that can bias inference and conservation decisions. Current approaches to address mislabelling rely on expert taxonomists validating every record. This approach makes intensive use of a scarce resource and reduces the role of the citizen scientist.
2. Species, however, are not confused at random. If two species appear more similar, it is probable they will be more easily confused than two highly distinctive species. Identification guides are intended to use these patterns to aid correct classification, but misclassifications still occur due to user-error and imperfect guidebook design. Statistical models should be able to exploit this non-randomness to learn confusion patterns from small validation datasets provided by expert taxonomists, yielding a much-needed reduction in expert workload. Here, we use a variety of Bayesian hierarchical models to probabilistically classify species based on the species-label provided by the citizen scientist. We also explore the utility of guidebooks provided by the citizen science schemes as a prior for species similarity, and hence draw conclusions for their future improvement.
3. We find that the species-label assigned to a record by a citizen scientist, even when incorrect, contains useful information about the true species-identity. The citizen scientists correctly identify the species in around 58% of records. Using models trained on only 10% of these records (validated by experts), we can correctly predict species-identity for 69 (90%CI: 64-73)% of records when the guidebook is used, vs 64 (58-69)% for models that do not use the guidebook. The fact that misclassifications can be predicted systematically indicates that improvements could be made to the guidebook to reduce misclassification.
4. By using Bayesian, hierarchical models we can greatly reduce the workload for experts by providing a probabilistic correction to citizen science records, rather than requiring manual review. This is increasingly important as the number of citizen science schemes grows and the relative number of taxonomists shrinks. By learning confusion patterns statistically, we open up future avenues of research to identify what causes these confusions and how to better address them.

## 2 Introduction

Citizen science has become increasingly important for addressing modern ecological questions. Systematic methods for monitoring biodiversity can rarely achieve the same scale in space or time. Simultaneously, these projects empower members of the public to take an active interest in the natural world and its stewardship [1]. However, these data are challenging to analyse, with heterogeneous recording effort across space, time, and taxa, and frequent misclassification of species [2]. A large number of models have been developed to tackle heterogeneous effort problems in citizen science data [3–5], however, fewer attempts have been made to address species misclassification [6, 7].

Misclassifications can affect biological inference. For example, in a survey aimed at describing the habitat associations of a focal species, mislabelling of other species as the focal species can increase both bias and uncertainty in the perceived habitat usage. If the habitat usage by the two species overlaps, this may be small, however, if species misclassifications lead to false positive records the resulting inference will be biased [8, 9]. Schemes that acknowledge these problems tend to tackle species misclassification via labour-intensive manual review of each record by experts [10]. Unfortunately, expert reviewers are few [11] and citizen science records are many [12]. We therefore require solutions to the problem that make efficient use of small amounts of expertly reviewed data.

Fortunately, most species are not confused at random [10]. Species are most likely to be confused if they share similar physical traits, such as coloration, size or behaviours. Such non-randomness is amenable to statistical treatment via the development of suitable observation models. Statistically modelling the observation process allows us to learn which species are confused and to make probabilistic reclassifications. These misclassification patterns can be learned from expert-validated data alone [13] but to reduce the workload for experts we can incorporate prior knowledge about species similarity. Crucially, we can incorporate prior knowledge about species similarity *from the perspective of the citizen scientists*. The features that make two species look similar to a citizen scientist, for whom species identification may be a new experience, are likely to differ from those used by experts.

Fortunately, we have expert knowledge on what citizen scientists see codified in identification guides. These guides are developed by scheme organisers and use simple, easy-to-learn features to help citizen scientists distinguish species. Scheme organisers often incorporate knowledge from citizen science workshops and previous schemes when designing guides [14]. This may lead to the traits used in different guides varying for the same species group, however, most guides are characterised by minimal structure (i.e. they do not use keys or only use very coarse keys) and the traits selected tend to be easy to find and distinguish without previous experience. Translating guidebooks into formal priors is challenging due to this lack of imposed structure, as it is unlikely that the traits are weighted equally and citizen scientists will develop heuristic hierarchies of traits and combinations thereof.

Overcoming this challenge and including guidebooks in misclassification models should also allow us to assess guidebook design. Assuming the species are distinguishable, if the models do not find signal in which species are confused (with or without the guidebook prior), then the guidebook (or some other form of training) is performing well and misclassifications are down to user error alone. If the models do find a signal in which species are confused, then there is structure in the misclassifications not currently addressed by the guidebooks. If the guidebook prior is uninformative, it is not capturing features that lead to confusions. This may arise from the citizen scientists using traits that are not included in the guidebook or making errors that do not correspond to conventional traits, for instance, labelling the record as a species with a similarly spelled name or some species being culturally more important, for example, if they are rare or invasive [15]. If the guidebook prior is informative, it is generating confusions by making species appear overly similar. This may arise from including too many traits and placing insufficient emphasis on the most informative traits.

In this paper, we present a series of observation models that address species misclassifications explicitly. These models include multiple approaches to incorporating guidebooks as priors (including not incorporating the guidebook). The prediction performance of these models is tested under cross-validation with different amounts of training data using both simulated and real-world data from the “Blooms for Bees” citizen science scheme [10]. We discuss the implications of the results for guidebook design and suggest extensions to these models.

## 3 Materials and Methods

### 3.1 Modelling Problem

For a data set of *N* records we consider two *N*-length vectors: the record-labels (the species label chosen by the citizen scientist), ***U***; and the record-identities (the species that a record truly belongs to), ***T***. The elements of both vectors take values from a set of *M* levels representing the possible species.

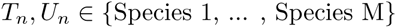

We construct our model generatively, beginning with the underlying biological process that determines the relative frequency of each species (the true record-identities). We assume this process is independent of our observation process, is proportionate to the species abundance and will often be the pattern we are interested in reconstructing from our data. Even if we are not interested in the biological process, by jointly modelling it with the observation process we can potentially improve our estimation of the species misclassifications [7].

In principle, any biological process model could be used. To simplify notation, we will assume the *m*th species’ relative frequency, *α*_*m*_, is a function of *K* environmental covariates (*X*, e.g. distribution of temperature, precipitation and wind): 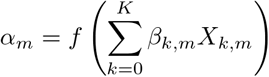.

Since the true record-identities, *T*_*n*_, are non-ordinal, the categorical (or single-trial multinomial) distribution, Cat, is a natural choice for the likelihood of any given record. The categorical distribution is parameterised with a vector of probabilities, ***A***, corresponding to the probability that the record truly belongs to each potential species. We therefore transform our unbounded linear predictor, ***α***, into a simplex using the softmax link function (a multivariate generalisation of the logit transformation, also known as the “multi-logit”).

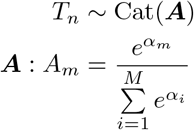

Now that we have a model for generating the true species identities, we need to link these identities to the labels assigned by the citizen scientists (i.e. the observation process model). We want to estimate the probability that a given record-label, *U*_*n*_, is generated conditional on the underlying record-identity, *T*_*n*_, Here, too, we will use a categorical distribution parameterised by the rows of an *M* × *M* matrix, ***C*** that corresponds to the record-identity, *T*_*n*_. This matrix encodes the pairwise confusability of *T*_*n*_ with each potential *U*_*n*_, i.e., the generation of *U*_*n*_ from *T*_*n*_. The formulation of 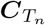 could take multiple forms depending on the amount of trust placed in the citizen scientist, whether external information is incorporated, and, if so, how that information is incorporated. Here, we describe the generic structure using the shorthand ***f*** for the different modelling frameworks, with ***f*** indicating the vector of classification probabilities conditional on the true identity of the record, and *f*_*m*_ showing the *m*^th^ element of that vector. Below, we describe the different variants of the modelling framework in more detail (see also Figure 1)

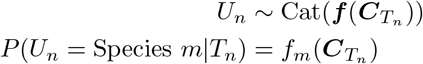

**Figure 1:**
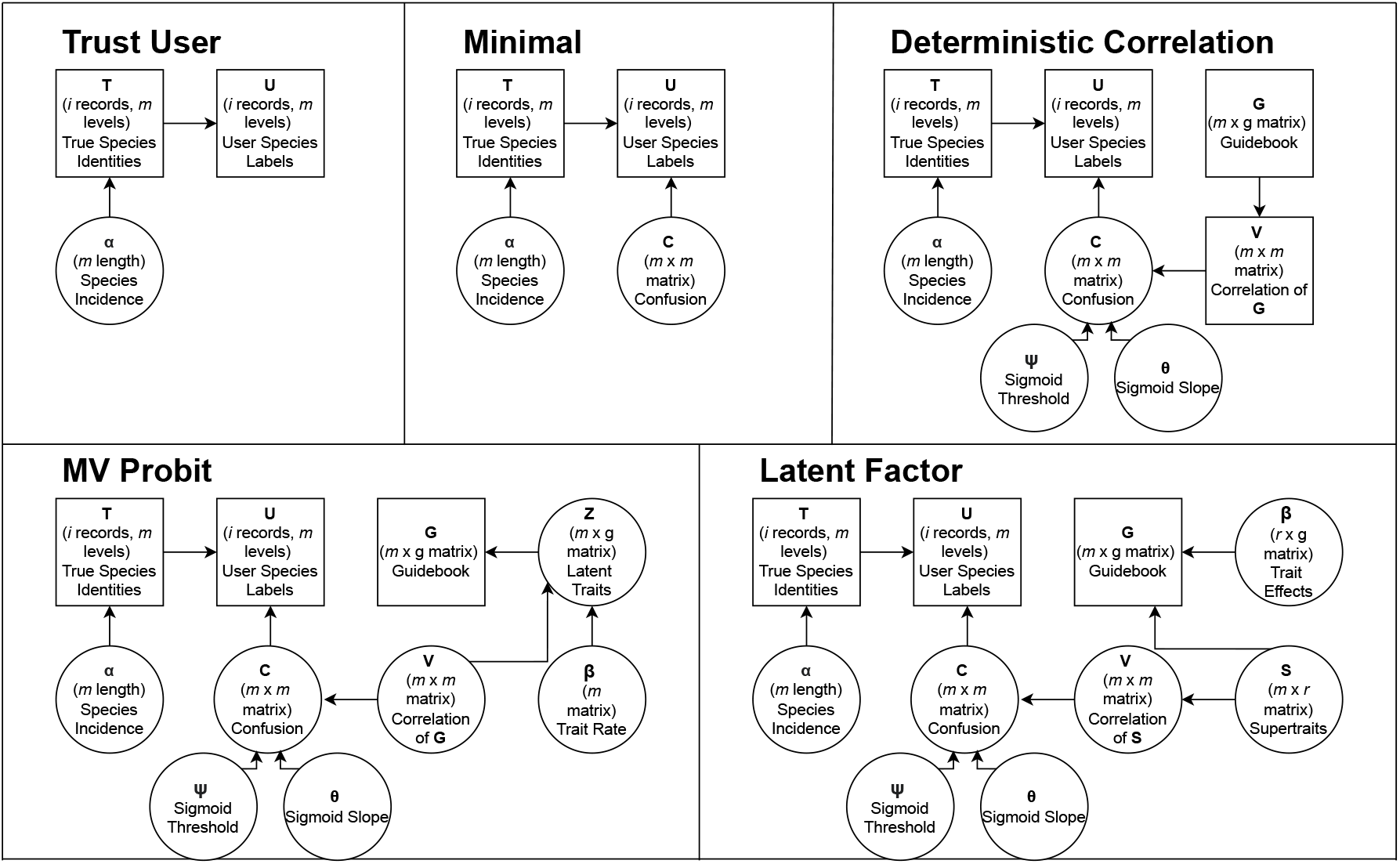
Causal Structures of Candidate Models. We define five modelling frameworks of different degrees of complexity. All of the frameworks contain a “Species Incidence” term, **α**, which corresponds to the function that allows the estimation of (and adjustment for) relative species abundance. The “Trust User” framework assumes that the record-label matches the record-identity. The “Minimal” framework incorporates an unstructured confusion matrix, **C**, allowing the record-label to differ from the record-identity. The “Deterministic Correlation”, “MV Probit” and “Latent Factor” frameworks all use the citizen science scheme’s guidebook, **G**, to estimate correlations between species, **V**, to inform **C**. The “Deterministic Correlation” uses the empirical correlation between the species in the guidebook as data to inform **C**. The “MV Probit” framework estimates the correlation between the species in the guidebook using a multivariate-probit model. These two approaches weight the guidebook traits equally. The final framework, the “Latent Factor” approach, is the most flexible, using latent factors, **S**, to combine and reweight the traits. All the approaches which use the guidebook are subject to a flexible sigmoidal transformation using a Normal CDF parameterised by θ and ψ

### 3.2 Model 0: Trust User

Our null “model” assumes that the citizen scientists are 100% correct in their labels (i.e. the label always matches the species identity). This model is extreme since even experts are rarely 100% correct but, implicitly, this is the model assumed by any analysis of citizen science scheme that uses the data without correction or mediation via calibration datasets and strong priors.

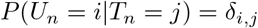

### 3.3 Model 1: Minimal

The simplest model (conceptually) is to populate the elements in ***C*** with parameters that allow the citizen scientists to confuse species (i.e. suggest the wrong label conditional on the species’ identity). If we do not know *a priori* which species the citizen scientist is likely to confuse, we can use free parameters to populate ***C***. The elements of ***C*** can be drawn independently and the rows used to parameterise the categorical distribution under softmax transformation. To make the model identifiable, we must fix one parameter in each row (i.e. in each vector passed to the categorical distribution). Here, we fix the correct classification (the diagonal of ***C***) to one to maintain consistency with our later models that use a correlation structure (and thus also have ones on the diagonal). As we expect correct classification to be more likely than any given misclassification, we centre the off diagonal values around zero. The spread of misclassification values is determined by a standard half-Normal prior on *σ*. The larger *σ* is, the greater the spread in probabilities of misclassification. A smaller *σ* indicates most misclassifications are equally likely to each other, but much less likely than correct classifications. It is plausible that *σ* could vary by species so we examined two versions of the model: one with a global *σ* parameter (as below) and a more flexible one where it is indexed by species, *σ*_*i*_.

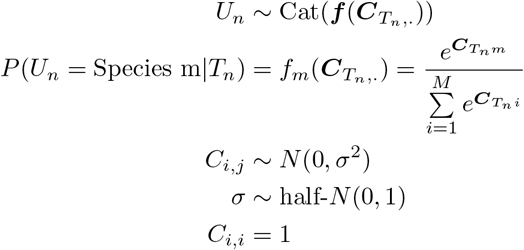

### 3.4 Estimating Species Similarity from ID Guides

Species identification by citizen scientists is done visually and we would expect similar-looking species to be more readily confused than highly distinctive-looking species. However, the question then arises: how do we know which species look similar to the citizen scientists? One source of information on this could be the identification guides provided to the citizen scientists by the scheme-organisers. These guides are often very different from professional taxonomic guides where subtle features and highly structured keys are relied upon. Citizen science guides are characterised by easy-to-recognise features and limited structure. Many guides are thus easily converted into simple *M* × *H* binary trait matrices, ***G***, with each row corresponding to a species, each column the level of a trait and a binary indicator in each cell indicating whether the indexed species has the indexed trait.

The distance between species in ***G***-space, ***V***, therefore, represents our prior expectation of which species are likely to be confused and can be used to inform our ***C*** matrix. There are multiple options for defining this distance depending on the amount of flexibility given to estimating the correlation in ***G***-space and the weighting of the different dimensions (i.e. the different traits). The simplest of these methods, the “Deterministic Correlation” model described below, is the only method to use the empirical correlation of species in ***G***-space as a measure of distance, making the least flexible use of the guidebook data. The “MV Probit” method relaxes this approach by estimating the correlation of species in ***G***-space as a measure of distance by means of a multivariate probit. Both of these methods apply equal weighting to the trait dimensions in ***G***-space, but in reality it is unlikely that all traits are equally important to the citizen scientists for species determination. For example, some traits, like colour, may be easier to assess without specialist knowledge and be relied on more heavily. They may also be viewed in combination, so while head colour, thorax colour and abdomen colour are separate traits in the guidebook, many species have the same colour on multiple body parts and will be thought of as “the ginger bee” (like many of the carder bee species). Our final method, the “Latent Factor” model, accounts for this behaviour by allowing the traits in ***G***-space to be up- or down-weighted and recombined using latent factors (or “supertraits”). The correlation between species in latent factor space is then used to inform ***C***.

The correlation distance between species, regardless of how it is estimated, does not necessarily map directly onto the confusability distance in the probability space. While we would generally expect the confusability ranking to be maintained, the scalar distance in the two spaces will likely differ. We, therefore, introduce a convolution step, wherein we apply a flexible sigmoidal transformation to each vector of correlational distances. Specifically, we use the Normal CDF function which has two parameters, a slope (*θ*) and threshold (*ψ*). As the slope approaches zero, the sigmoid becomes a step function, with values smaller than the threshold transformed to zero and larger values become ones. The threshold determines where this step occurs. As the slope gets larger, the values around the threshold will become values between zero and one, with only more extreme values becoming zeroes or ones. We expect a small number of species to be confused with the true species, so place a prior having a mid-high threshold and small slope. Similarly to the *σ* parameter in the “Minimal” model, it is plausible that *θ* and *ψ* could vary by species so we examined two versions of the model: one with global values for those parameters parameter (as below) and a more flexible one where they are indexed by species (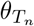 and 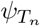).

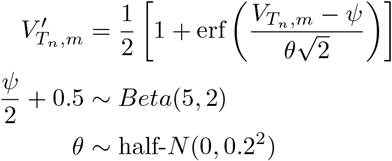

We also need to allow for confusion not associated with the guidebook. We achieve this by incorporating 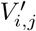 as a prior for *C*_*i,j*_. As 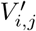 is bounded by zero and one, we use a Beta distribution with a mean-variance parameterisation. This parameterisation allows the variance, *λ*, to up or down-weight the contribution of 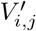 to *C*_*i,j*_.

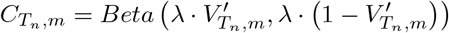

Although *C*_*i,j*_ is bound by zero and one, the vector still needs to be normalised to generate a simplex of confusions. One option would be to use the softmax transformation (as in the Minimal Model), however, with constrained values (unlike in the Minimal Model) softmax tends to generate a large number of small probabilities. This contradicts our understanding of confusions which we expect to be sparse, with a few large probabilities (commonly confused species) and many zero probabilities (species which are never confused). Fortunately, as the values are now all positive, we can simply normalise by dividing each element of the vector by the sum of the vector. This is easy to calculate and is compatible with sparse probabilities.

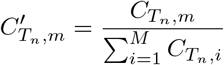

These normalised values can now be used to parameterise the categorical distribution as in the Minimal Model:

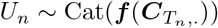

### 3.5 Model 2: Deterministic Correlation

There are several numerical methods for calculating the empirical correlation of a dataset. These point estimates for correlations, e.g. Pearson’s, Kendall’s and Spearman’s coefficients assume no uncertainty in the correlations but are very computationally efficient, and thus useful as a baseline guidebook-based model. We use the Pearson’s correlation coefficients for the species in the guidebook as data, ***V***.

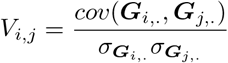

### 3.6 Model 3: Multivariate Probit

The next most complex version of the model allows the uncertainty in the guide-based correlation, ***V***, to be estimated. Since we have binary data, a natural method for doing this is the multivariate probit, which assumes that binary variables are realisations of correlated, normally distributed latent processes [16, 17]. This assumption is both computationally expedient and often matches our biological knowledge, since many binary variables are functions of continuous underlying processes. For example, an individual is considered infected or not infected (binary) based on an underlying quantity of pathogens (continuous).

The binary data, ***G*** formed of *H* traits and *M* columns, are linked to the latent continuous variable, ***z***, by means of a thresholding function, 𝕀, which returns a one if the latent variable is positive and a zero if it is negative. This thresholding process is equivalent to a probit link.

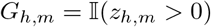

The latent state, ***z***, is generated from an *M* length intercept-only linear predictor, ***β*** and a multivariate-Normal (MVN) error term, ***ϵ***. The correlation between species is induced through the normalised covariance matrix parameter of the MVN.

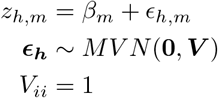

The intercept corresponds to the number of traits each species has (i.e. the number of traits indicated by a 1 in the guide). There is little biological interpretation for this value (as it is determined by encoding decisions), so we use a standard Normal prior which is minimally informative under a probit transformation.

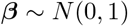

We place an LKJ prior on the correlation prior with *η* = 1. The LKJ prior corresponds to beta-distributed marginal correlations of Beta 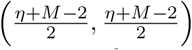. At *η* = 1 this is relatively uniform at small values of *M* but with slight peaking at 0 correlations for larger values of *M*. Lower marginal correlation between species as they increase is plausible although the degree of shrinkage should be monitored when re-applying. There are relatively few priors for correlation matrices and the LKJ distribution is computationally efficient.

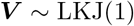

### 3.7 Model 4: Latent Factor Models

The approaches thus far all treat traits as equally important, however, this is unlikely to hold in reality. Field biologists (particularly ornithologists) have long referred to the “jizz” or “vibe” of an organism: the combination of shape, mode of movement, posture, colouration and myriad subtle traits which allow an organism to be identified from a quick glance. In these scenarios identification is not occurring on a trait-by-trait basis but some reading of the whole or of groups of traits together. While non-expert users may not achieve identification at a glance, it is likely they process the guidebooks and identification process in a similar way. Some species will be discounted immediately based on the dominant colour (a combination of thorax, abdomen, and tail colour), for instance, while other species will be more challenging to disentangle. The development of citizen science identification guides is often an attempt to formalise this process by using measurable traits. Guides may be able to capture this with expert construction, but the need to function for novice citizen scientists as well as more experienced observers means that there will often need to be an imperfect match between the guide layout and its use by groups of different experience levels. As a result, to understand guide-based confusion we need a system by which we can up- and down-weight the contribution of different traits in the guidebook.

One approach is to consider the traits in ***G*** as functions of latent factors, ***S***. These latent factors can be thought of as “super-traits”: continuous underlying processes that when combined in different proportions give rise to the traits that are measured and included in the guide. Crucially, the correlation between latent factors represent the similarity between species as seen by the citizen scientists, so correlations between species in ***S*** give us ***V***. The number of latent factors, *R*, can be estimated or may be pre-determined based on the size of data/previous experiments (or the latter used to inform a prior for estimation).

#### Estimating Latent Factors from Traits

Most traits are binary so here we link traits, ***G***, and latent factors, ***S***, using logistic regression. The latent factors need to be flexible but estimable. For this reason, we assume a linear relationship between traits and their latent factors and provide standard Normal priors which are relatively uninformative under logit transformation. Where more information is known about the latent factors, more complex functional forms could be used. Similarly, where non-binary traits are present the approach can be generalised to accommodate more complex traits using other GLM formulations.

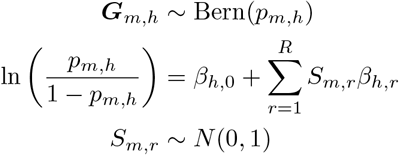

As the latent factors have exchangeable priors, we risk label-switching identifiability issues (i.e. the indexing of the latent factors may be inconsistent between MCMC chains). We need to impose some form of order on ***S***, however, as we are going to estimate the correlations between the rows of ***S*** (i.e. between the species in super-trait space) ordering the latent factors is undesirable. For this reason, we place a hierarchical prior on ***β***.

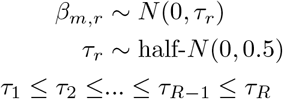

Even with these restrictions, ***S*** and ***β*** are only identifiable when linked to another data source (in our case, the identity-label confusions).

#### Linking Latent Factors to Species Confusions through

***V*** To link ***S*** to ***V*** we simply calculate the correlations between the rows of***S*** (i.e. between the species in latent factor space). To do this, we must first z-score normalise the rows of ***S*** to achieve a mean of zero and variance of 1 to give us 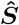. We can then calculate the Pearson correlation of ***S*** by post-multiplying 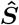 by it’s transpose to give us ***V***.

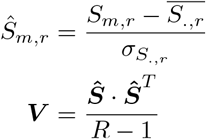

### 3.8 Measuring Performance

The aim of these models is to reduce the work required by expert taxonomist reviewers to processing a small validation data set to which the model can be fit, while propagating and stating the uncertainty in corrected classifications. The best model, therefore, is the one that has highest out-of-sample predictive power from the smallest training data set. In this section, we define how we measure out-of-sample predictive performance using the correct classification rate and our experimental design for comparing performance across different sample sizes.

First, since the same performance metrics are measured across varying models we will use the following summary notation to represent all the models (including those that incorporate ***V***):

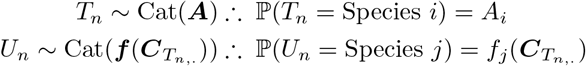

If we then want to predict *T*_*n*_ given *U*_*n*_ we apply Bayes rule:

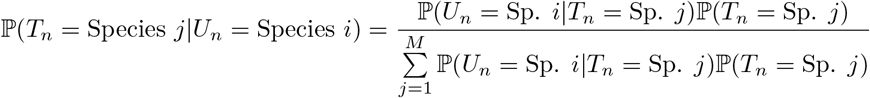

We will represent the *M* × *M* matrix that defines all the possible combinations of *U* and *T* using the symbol **Ψ**. Each row of **Ψ** corresponds to a label and defines a simplex (probabilities summing to 1) which give the probability of the possible record-identities, *T*.

The simplest way to think about model performance is to measure how often the record-identity predicted by the model matches the true record-identity as validated by the model, i.e. the correct prediction rate, R. To estimate this value, we generate prediction values, 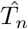, for each record in a holdout set of size *N*, and measure the proportion of records for which 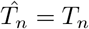

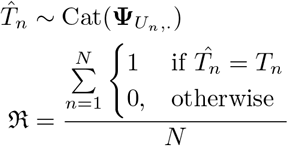

### 3.9 Comparing Performance under Cross-Validation and Varying Data Richness

As outlined above, models need to be evaluated on both predictive performance under varying data richness (i.e., what proportion of the available data is used in training the model). We therefore use holdout cross-validation, where the data available are randomly assigned to two groups, *d*_0_ (training) and *d*_1_ (testing). The assignment is repeated *J*-times to estimate the average and spread of the performance metrics. Varying data richness (the sizes of *d*_0_ and *d*_1_) introduces two sources of uncertainty associated with the training and testing sets of data. As either the training or testing set shrinks, the number of data-point combinations will increase, leading to higher variability in the fitting process and prediction targets that will both be reflected in the performance metrics.

We are not interested in uncertainty due to testing set size therefore the simplest solution to this source of variation is to fix the size of *d*_1_. Naturally, the size of available testing data is complementary to the amount of available training data. We therefore chose to test on 25% of the available data, as requiring more than 75% of the available data to be used in training would not represent a significant reduction in the work of the expert validators.

Varying the size of *d*_0_ introduces two sources of variability. Firstly, there is the larger uncertainty in parameter estimates associated with smaller data sizes. Secondly, there is the larger number of combinations of data that may be used in the training data. The first is of vital importance to understanding model performance while the second is a nuisance that we should control for. Unfortunately, it is hard to predict what impact the latter will have, so we have to take a computationally-intensive approach.

We start by choosing a large value of *J* and running holdout cross-validation for all the model classes at the smallest *d*_0_ of interest. The smallest size of *d*_0_ will have the largest performance-metric variability due to training set effects (fortunately, they will also be the quickest models to run). We then repeatedly sub-sample from 1 to *J* of the cross-validation exercises and assess at what value, *J* ^*′*^, the centre and spread of the performance-metrics stabilises. The larger sizes of *d*_0_ can then be run only *J* ^*′*^ times and the variability therein can be attributed solely to uncertainty in parameter estimation.

### 3.10 Case Study

We apply our modelling framework to real-world data from the “Blooms for Bees” citizen science program. In this program, citizen scientists were asked to photograph and identify to species-level every visiting bumblebee to a single plant with at least one open flower in their garden or allotment. The scheme provided an unstructured identification guide to participants via a mobile phone app (through which they also submitted their records). The guide has 23 bumblebee species and 69 traits (including levels of traits). Falk *et al* then reviewed the photographs and corrected any misclassified species [10]. This generated 2314 records containing the original label from the citizen scientist and the corrected identity provided by the expert reviewer. The records are primarily concentrated around the West Midlands of England as the program was developed by Coventry University. This restricted geographic region, and focus on garden and allotment habitats allowed us to adopt a very simple biological process model using an intercept-only linear predictor for each species, ***α***, corresponding to their relative abundance. The performance of the observation models is compared using the cross-validation protocol above.

### 3.11 Simulation Study

In some instances, we may not know exactly which guidebook the citizen scientists used. For example, participants may supplement the guidebook provided by the scheme with their own favourite guidebook. It is therefore important to understand how sensitive our models are to the exact guidebook used. We can explore this using simulations. Guidebooks for the same group of species may differ in a large number of ways - the rank order of species similarities, the frequency of the traits used, the determination of the correlations between species (i.e., how strongly correlated the species are in the trait-space defined by the guidebook). To assess the sensitivity of the models to these changes, we need to simulate under one guidebook scenario and then compare prediction performance when the model is fit with the correct guidebook vs a contrasting one. These kinds of transplant tests are computationally intensive, especially when testing under the cross-validation conditions described above.

To make these simulations computationally tractable, we need to prioritise how we simulate and change the guidebook. First, we choose one of our observation models to be the basis of the simulation. The “Multivariate Probit” model is the simplest model that allows us to generate a full guidebook. In this model (Figure 1), we can change the rank ordering of species similarities by re-organising the columns of the correlation matrix, the frequency of the traits using the mean *β* parameter, and the determinant of correlations by modifying the prior on ***V***. This brings us to the second prioritisation: which of these to change. The guidebook-space defined by these parameters is huge and not practical to explore fully. We choose to focus only on changing the rank order of species similarities as this is a commonly discussed decision by guidebook designers (e.g., when navigating a dichotomous key designers often try to ensure the final pair of species in each branch are easy to distinguish). Finally, as the “Multivariate Probit” model is stochastic, we need to repeat simulations to account for the inherent noise in the generative process. For this reason, we limit our comparison to two contrasting scenarios (i.e., two species rankings) to facilitate more repetitions.

In order to make our simulations realistic, we draw the parameters for each scenario from the posterior of the “Multivariate Probit” fit to the real data. To change the species rankings in a consistent way, we modify the correlation parameter using hierarchical clustering with the complete linkage algorithm implemented using the “hclust” function in the R “stats” package [18, 19]. The precise nature of the reordering is not significant, but it is worth noting that by only reordering the correlation matrix we keep the same matrix determinant.

We now have full parameters for two contrasting scenarios (Figure 2). The “Real” scenario has the same rank species similarity as our real data while the “Restructured” scenario has a different rank species similarity

**Figure 2:**
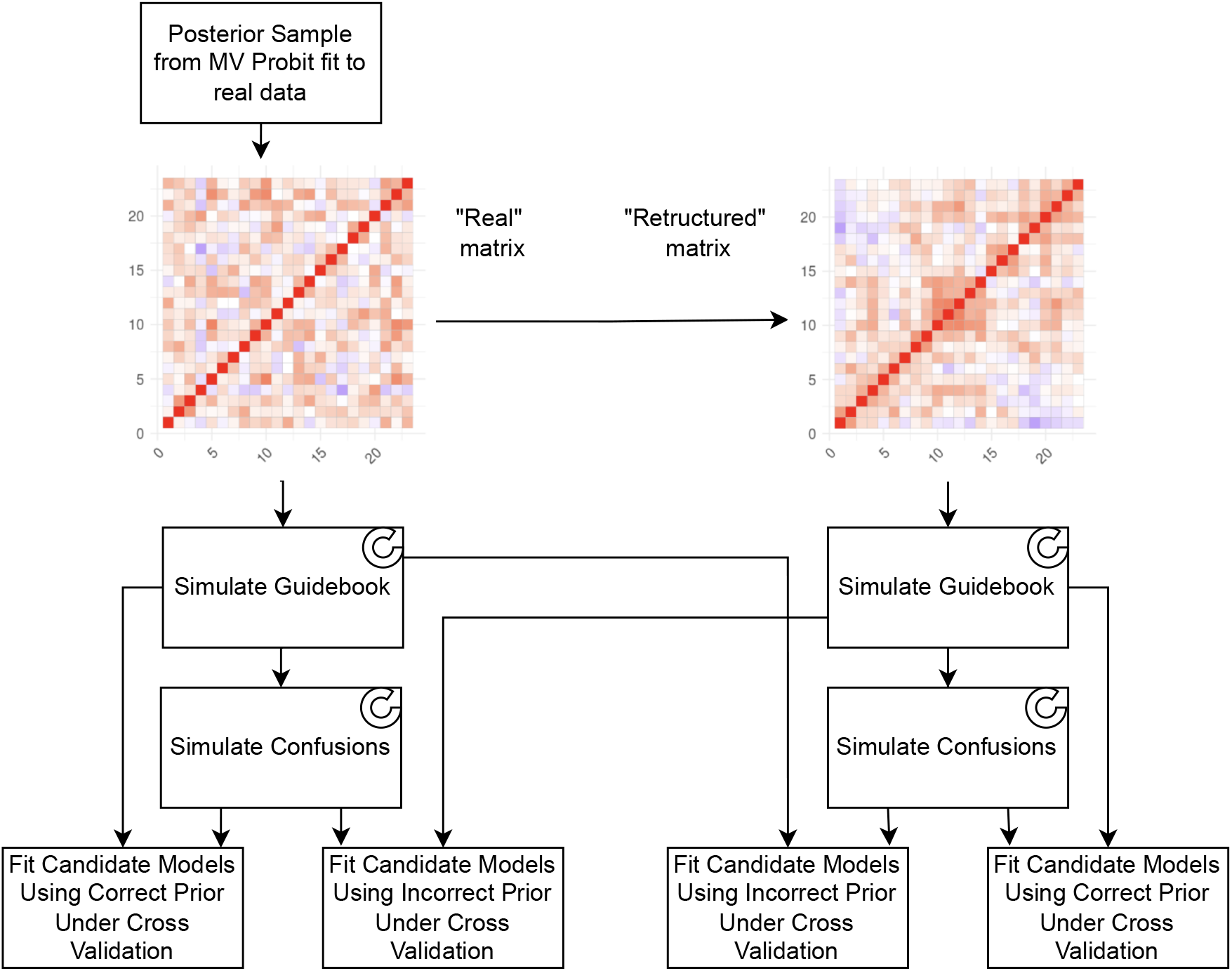
Simulation study outline. The simulation scenarios are based on species correlations estimated using the real data. This correlation matrix (“real”, with a blue to red scale indicating negative to positive correlations) is used to generate one set of simulations directly, and restructured to generate a contrasting correlation matrix (“restructured”) from which distinct but comparable simulations are created. Each of our candidate models is then fit (under cross validation) to the simulated data using either the correct or contrasting guidebook as a prior. This allows us to assess how sensitive to the guidebook the models are. Steps which are repeated are indicated with a partial concentric ellipse.

We take five samples from the posterior of the “Multivariate Probit” model fit to the real data. From these samples, we generate five guidebooks and five corresponding data sets (i.e., vectors of species labels and species identitities). To these data, we fit 7 candidate models (Minimal, Deterministic Correlation with correct and incorrect prior, Multivariate Probit with correct and incorrect prior and Latent Factor with correct and incorrect prior) under the same cross-validation scheme described for the real data.

## 4 Results

### 4.1 Computational Resources

Analyses were run in R (v4.2.1)[18] with the CmdStanR (v0.5.3)[20] interface to Stan (v2.30.1)[21] on a 64-bit workstation with 32 AMD Ryzen Threadripper 3970X CPUs running a Ubuntu 20.04.5 LTS operating system.

### 4.2 Model Convergence

All models achieved convergence with 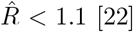, *<* 2% divergent transitions [23], and effective sample sizes of over 100 samples/chain (with the exception of a small number of lower level parameters in the “Latent Factor” model) [24]. Model runtimes varied by model type and data richness, with the more complex models and larger datasets taking longer to run (see Figure 3) with runtimes ranging from a minute to just over 3 hours.

**Figure 3:**
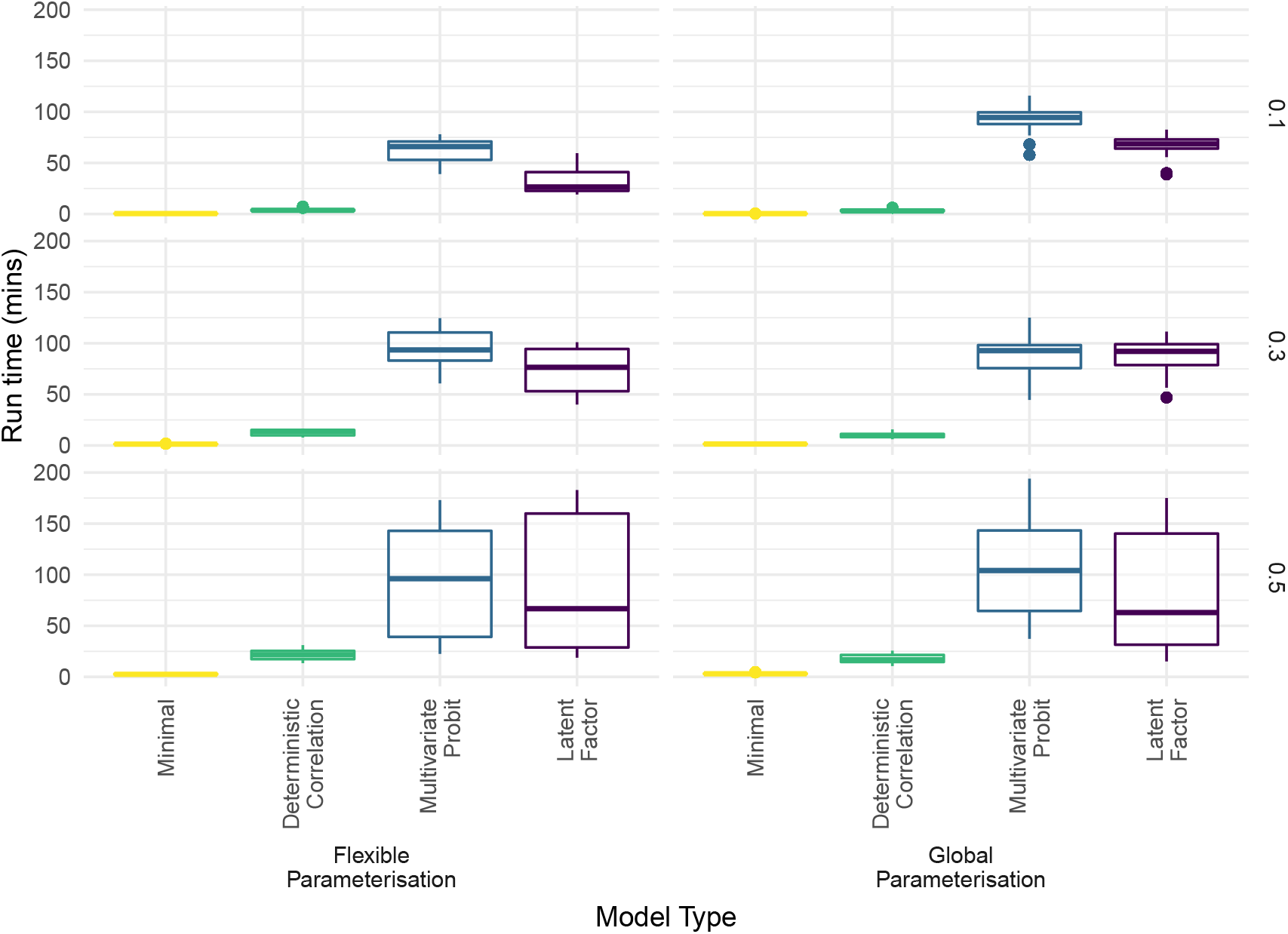
Comparison of model run times. Median and interquartile ranges for run times across cross-validation model fitting. Each row panel corresponds to different levels of data richness. As model complexity and data richness increases, models take longer. Run times vary from minutes to a few hours.

### 4.3 Case Study

When tested against the real data at the lowest level of data richness (10%), the best median performance of models which used the guidebook exceeded the “Trust User” model by around 10% and the “Minimal” models by 5%. The best performing parameterisation of the “Minimal” model predicted the proportion of correct classifications was 0.64 (90% Credible Interval (CI): 0.58-0.69), compared to the best performing parameterisations of the “Deterministic Correlation”, 0.67 (0.62-0.71), the “Multivariate Probit”, 0.69 (0.64-0.73), and “Latent Factor”, 0.69 (0.64-0.73) models.

With higher data richness, the differences in model performance shrink and performance quickly plateaus. Indeed, between data richnesses of 30% and 50%, the best performing parameterisations of the models have the same median performance, with only small improvements in precision (if any). The “Minimal” model achieves a correct classification rate of 0.68 (0.64-0.71) at 30% with the 90% CI shrinking to 0.65-0.71 at 50%. These rates are very close to those achieved by the guide-based models across which there is almost no difference in performance. The “Deterministic Correlation” model achieves the same rate of 0.7 (0.67-0.73) at 30% and 50% data richness. The “Multivariate Probit” yields 0.7 (0.66-0.73) at 30% with the credible interval shrinking slightly to 0.66-0.72 at 50% data richness. The “Latent Factor” performs identically at both levels of data richness, with rates of 0.69 (0.66-0.72).

The flexibility of the parameterisations tested generally made little difference in performance except at low data richness. For each model, we tested two parameterisations and compared their 50% credible intervals (more sensitive to differences than the more conservative 90% CIs used for between-model comparisons). As shown in Figure 4, the less flexible parameterisation of the “Minimal” model performed best at 10% data richness (0.64 (0.62-0.66) vs 0.62 (0.6-0.64)), with no difference at higher levels of data richness. In contrast, the more flexible parameterisation of the “Deterministic Correlation” model performed slightly better at all levels of data richness. The “Multivariate Probit” and “Latent Factor” models seemed less affected by parameterisation (although the less flexible parameterisation of the “Latent Factor” model had much larger uncertainty in the tails at 10% data richness than the corresponding flexible parameterisation). Based on these results, the best model fits to the simulated data are the less flexible parameterisation of the “Minimal” model and the flexible parameterisation of the other models.

**Figure 4:**
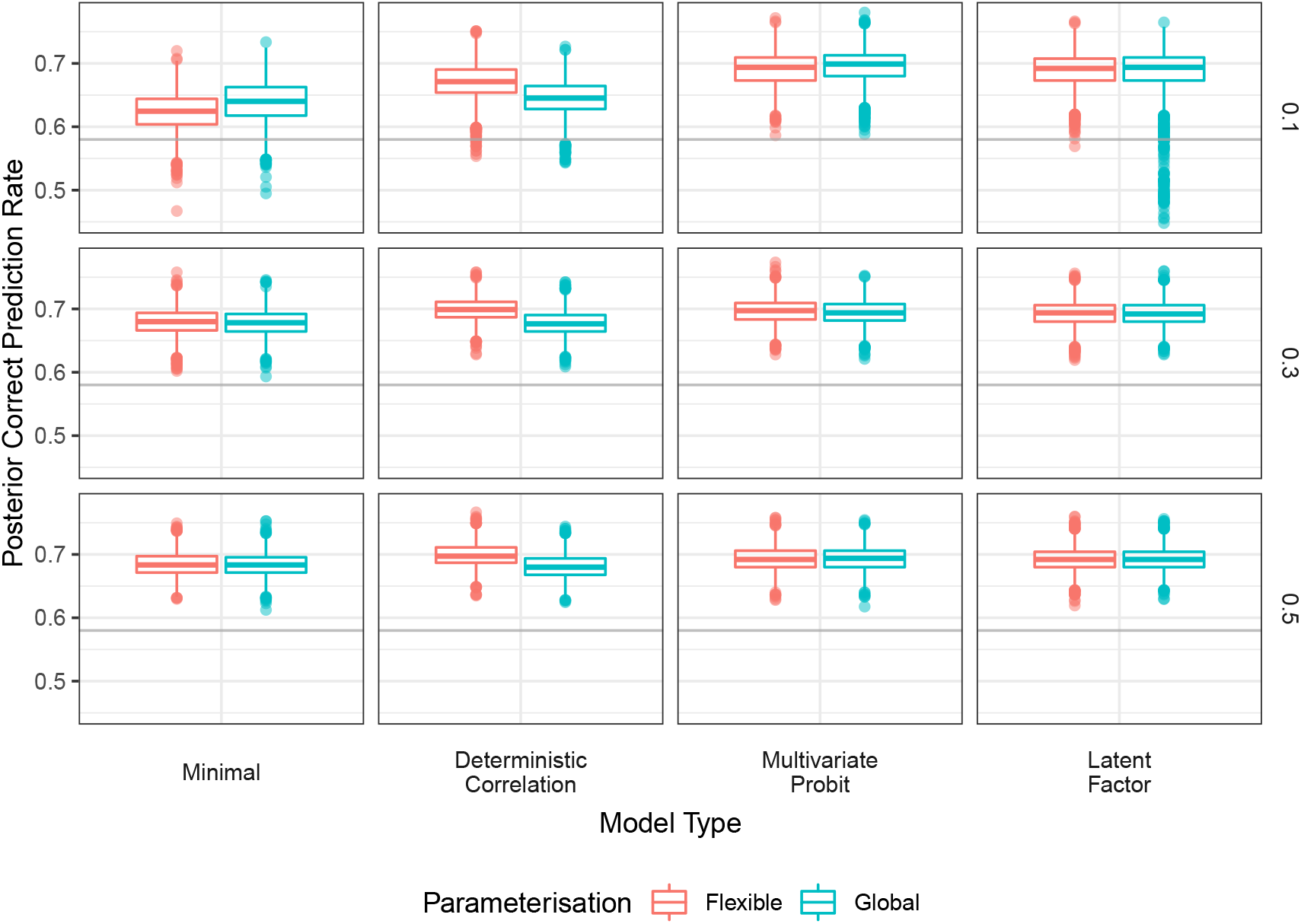
Comparative performance of models fit to real data. Aggregated posterior correct classification rate for models fit under cross-validation to the real data. The data richness for the cross validation scheme is indicated by the horizontal panels (0.1=10% data richness, 0.3=30%, 0.5=50%) and model types by the vertical panels. There are two parameterisations for each model, the “flexible” one which allows the model to vary on a species-wise basis (the variance in the “Minimal” and sigmoidal transformation parameters for the others) vs “global” wherein these parameters are shared across species. The correct classification rate achieved by the citizen scientists is shown as a horizontal grey line.

### 4.4 Simulation Study

The difference in performance between the “Minimal” model and guide-based models was much greater in the simulation study at low data richness. At 10% data richness, the “Minimal” model achieved 0.23 (0.17-0.36) for the “real” simulations and 0.31 (0.19-0.4) for the “restructured” simulations, rates approximately 0.2 lower than the guide-based models. There is a consistent but small improvement of performance when the “correct” guidebook prior is used for the “Deterministic Correlation” and “Multivariate Probit” models, while the “Latent Factor” model performs equally well with either prior, Figure 5. For both simulation scenarios, the “Deterministic Correlation” receives a bump as a consequence of using the correct prior in median correct prediction rate of 0.02 and the “Multivariate Probit” one of 0.01.

**Figure 5:**
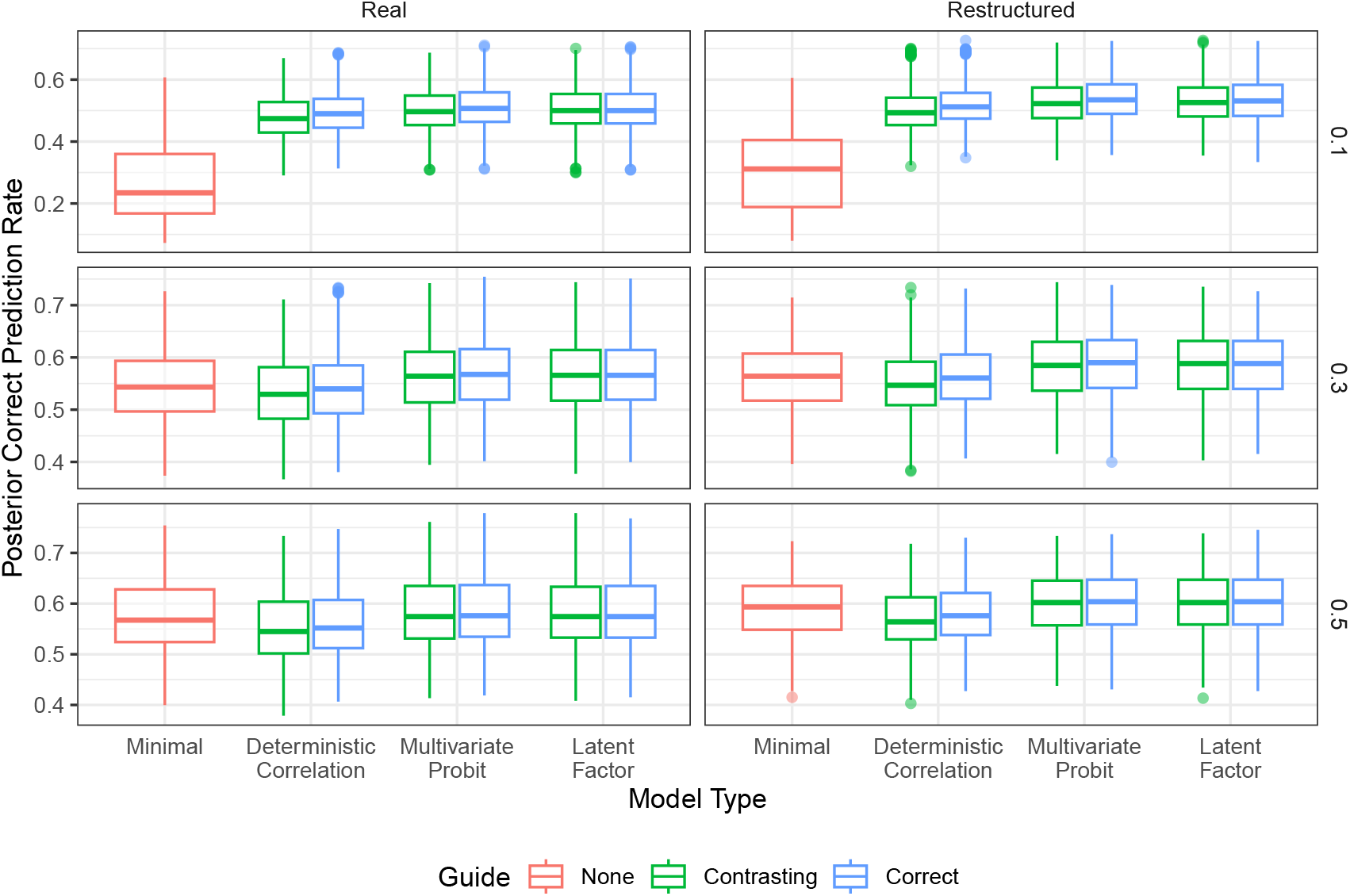
Comparative performance of models fit to simulated data. Data were simulated under two guidebook scenarios (indicated by the column titles), one drawn from the real data (“Real”) and one from a clustered adaptation (“Restructured”). Each model was then fit to each scenario and given either the matching (“Correct”) guidebook or the alternative (“Contrasting”) guidebook. The “Minimal” model does not use the guidebook prior. The data richness for the cross validation scheme is indicated by the horizontal panels (0.1=10% data richness, 0.3=30%, 0.5=50%).

At medium data richness, the results are similar for both sets of simulations, so henceforth for clarity the results reported are for the “restructured” simulation. At 30% data richness, the “Minimal” model achieves a rate of 0.56 (0.52-0.61), a slightly better performance than the “Deterministic Correlation” model with the contrasting prior, 0.55 (0.51-0.59), and the same as it with the correct prior, 0.56 (0.52-0.61). The “Multivariate Probit” model performs better with both the incorrect prior, 0.58 (0.54-0.63) and with the correct one, 0.59 (0.54-0.63). The “Latent Factor” model equals the latter performance with both prior types.

At the highest data richness, 50%, the “Minimal” model performance, 0.59 (0.55-0.63), exceeds that of the “Deterministic Correlation” model: 0.56 (0.53-0.61) with the incorrect prior and 0.58 (0.54-0.62) with the correct one. The “Multivariate Probit” and “Latent Factor” models now perform identically with either prior, yielding a correct classification rate of 0.6 (0.56-0.65).

## 5 Discussion

We have demonstrated that there are predictable patterns to how citizen scientists confuse species and these patterns are informed by the guidebooks they use. The current approach to correcting citizen science data requires labour-intensive expert review of every single record. We have shown on real data that it is possible to reduce this work by 90 percentage points yet maintain a high rate of accurate classifications (70% with the guidebook and 65% without) with appropriate probabilistic uncertainty for each classification (compared to 58% without uncertainty when you trust the citizen scientists).

Misclassifications that are, in part, predictable, indicate that the identification guides could be improved to exploit the structure in these confusions. If the guidebooks do not improve misclassification prediction, this would indicate that there are similarities between species that are not being captured by the guidebook. If the guidebook does improve misclassification prediction (as we found here), it means the guidebooks may be the source of the confusion, for example, by making species seem too similar. In principle, the informativeness of the guidebook could vary between species, however, the global parameterisation (which links the guidebook to species more equally) generally performed similarly enough to the more flexible parameterisations of the same model. That all the models reach the same maximum correct classification rate (approximately 70%) indicates that there is perhaps no pattern in the rest of the misclassifications, with the species being confused at random. These unstructured confusions are unlikely to be affected by improvements to the guide but could potentially be mediated by increased training of participating citizen scientists if possible.

When comparing these models, we are most interested in which performs the best with the least amount of validation data. In both the real data case and simulation study, the “Multivariate Probit” and “Latent Factor” models comfortably achieve the highest correct classification rate. The other models do improve with increased data and eventually match the performance of the others. For the “Minimal” model, this is likely because it relies on uninformative priors. The “Deterministic Correlation” model is fundamentally a point estimate version of the “Multivariate Probit” model (in terms of how they treat the guidebook). Essentially, this comes down to how estimable the correlation matrix is from the guidebook data. If the guidebook had a huge number of traits relative to the number of species, the two would give identical answers. In real applications, this is unlikely to ever happen (and with binary traits, correlations are even harder to estimate).

Our simulation study shows that using the same guidebook as a prior and to generate the data leads to a modest gain in performance for the guide-based models. The “correct” guidebook led to slightly improved performance of the “Deterministic Correlation” and “Multivariate Probit” models, and the “Latent Factor” model performed identically with the “correct” and “contrasting” priors. Nevertheless, at low data richness, these models all outperformed the “Minimal” models that used no guidebook. That the gains in model performance were only modest motivates a more thorough exploration of guidebook space. It’s notable that the “Latent Factor” model (where the correlational structure is between supertraits, rather than straight guidebook traits), shows no difference at all between the two priors. Understanding the interplay between these features should form the basis of future work if we wish to use these models to directly inform future guidebook design.

There are also several exciting extensions to these models that could be developed. Currently, the models are self-contained, relying on no collection of covariates or other data sources, but it would be possible to incorporate additional information in both the biological and observation components of the model. In our studies here, we have assumed an intercept-only model for species incidence. This could be extended to include more sophisticated models that account for species-habitat associations [25] or species interactions [26]. In modelling these processes jointly, there would be a feedback loop between the observation and biological process, improving both simultaneously, and propagating uncertainty. It is worth noting that species misclassifications is only one of several issues that need to be addressed in citizen science data [2, 3].

The misclassification component of the model could also incorporate covariates. For example, in an open meadow it might be easy to follow a bee until an identification can be made confidently, while more difficult terrain might make this impossible. We could also incorporate information about the observers themselves, such as their level of experience or track record of making correct identifications [27]. The existing model structures could be easily adapted to include these features by placing a linear predictor on ***C*** with ***V*** acting as a prior for the intercept (i.e. baseline confuseability).

The widespread use of mobile applications for data collection and submission opens up the possibility of deploying these data in real time [28–30]. Several citizen science apps already offer suggestions of potential similar species (sometimes weighted probabilistically) [31, 32]. These models could be used to suggest such alternatives and to feedback common confusions to the scheme designers. In turn, this could facilitate experimentation on guidebook design, where different users are given different versions of the guidebook, or the guidebook is adapted and updated live.

## 6 Conclusions

Modelling observation processes is a challenging but essential step in modern ecological research. Frequently, we must learn these processes directly from the data but here we have shown that there are useful priors available in the form of guidebooks. The mutual benefit of combining explicitly modelling the observation process with input from citizen science scheme developers is currently underexplored, particularly when it comes to misclassification. Leveraging statistical models can help reduce the workload of taxonomic experts and thus unlock the scalability of citizen science data for ecological research. The development of these methods relies on citizen science scheme organisers adopting positive attitudes to data sharing, and methods developers engaging positively with that community to learn from them.

## 7 Acknowledgements

We benefited greatly from discussions with Falk *et al* [10] and from their generous attitude to data sharing.

